## Supplementary material for "Structure-guided secretome analysis of gall-forming microbes offers insights into effector diversity and evolution": Fig. S1

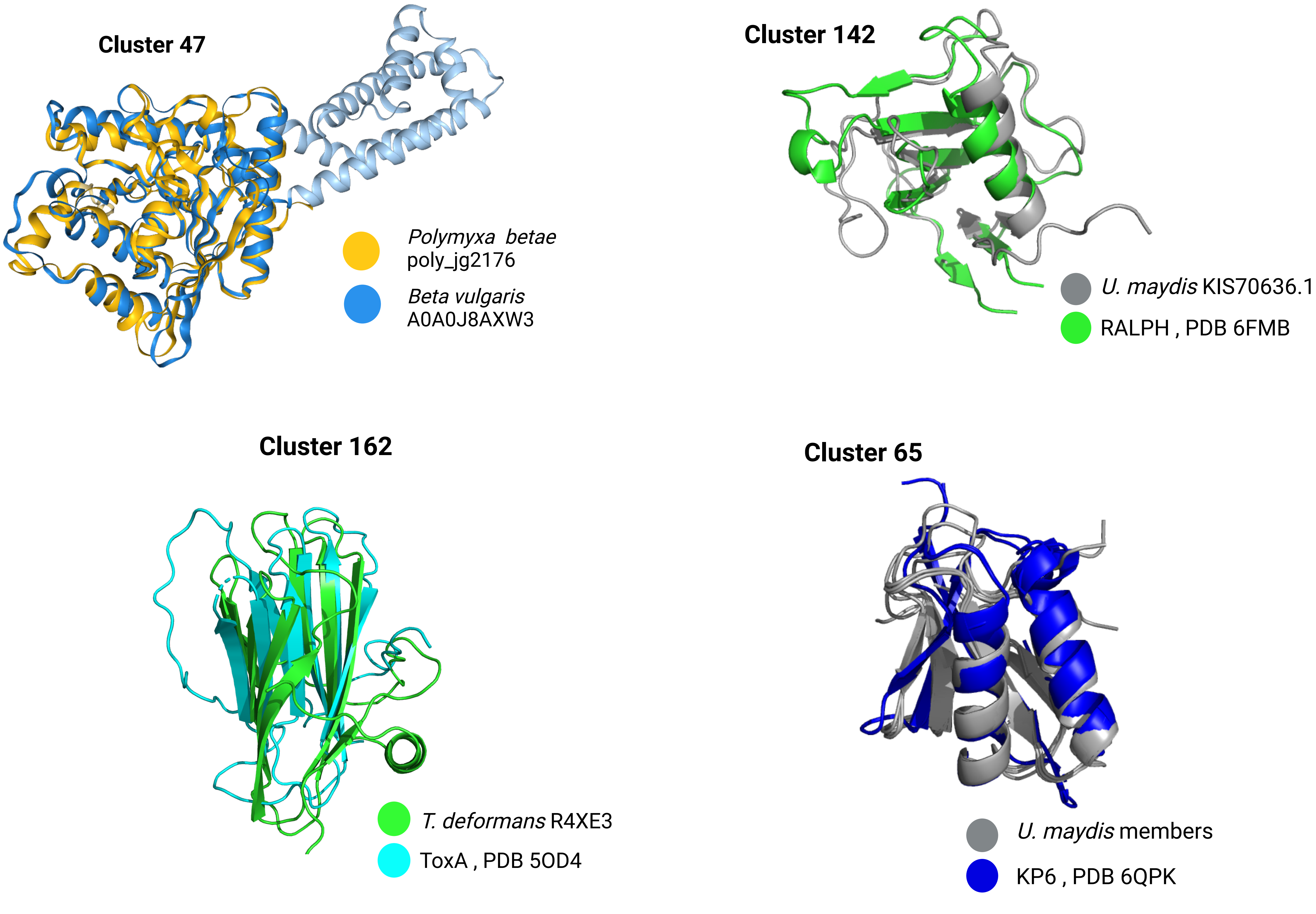


Structural alignment with the known effector families. The *Polymyxa betae* alignment with its host GTPase has been downloaded from the FoldSeek webserver.
