## Supplementary figures and images for "Structure-guided secretome analysis of gall-forming microbes offers insights into effector diversity and evolution"

### Fig. S2

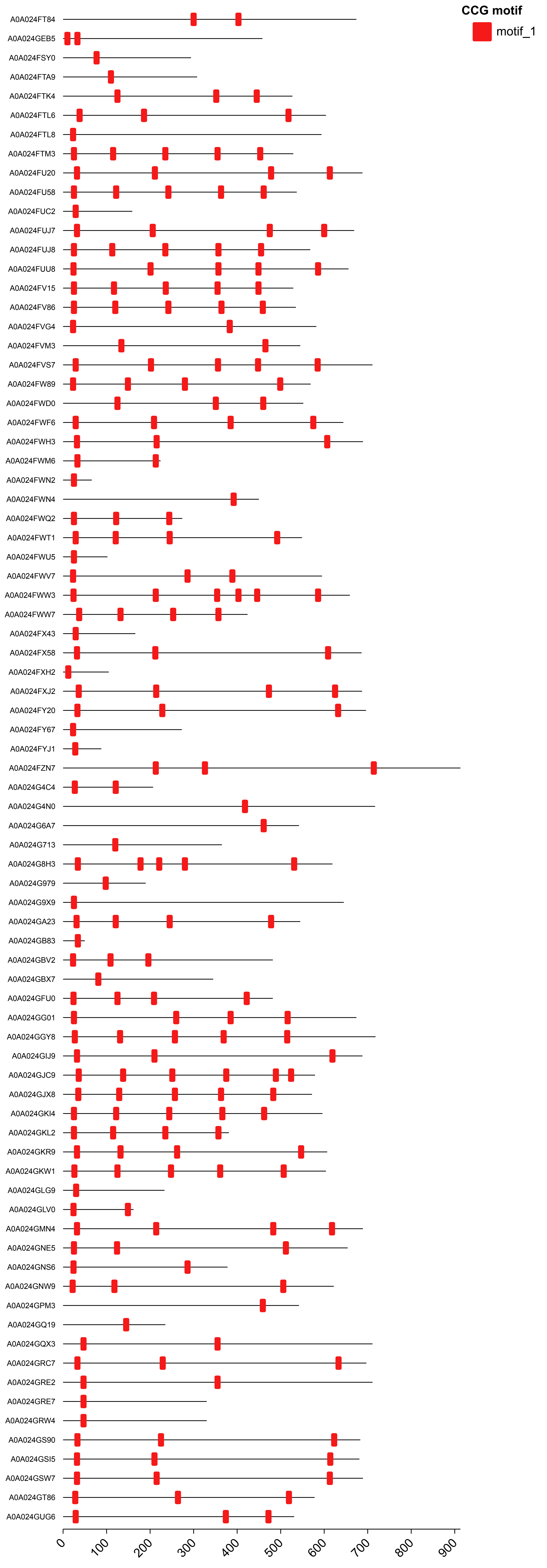


Location of CCG motifs identified by MEME scan

### Fig. S3

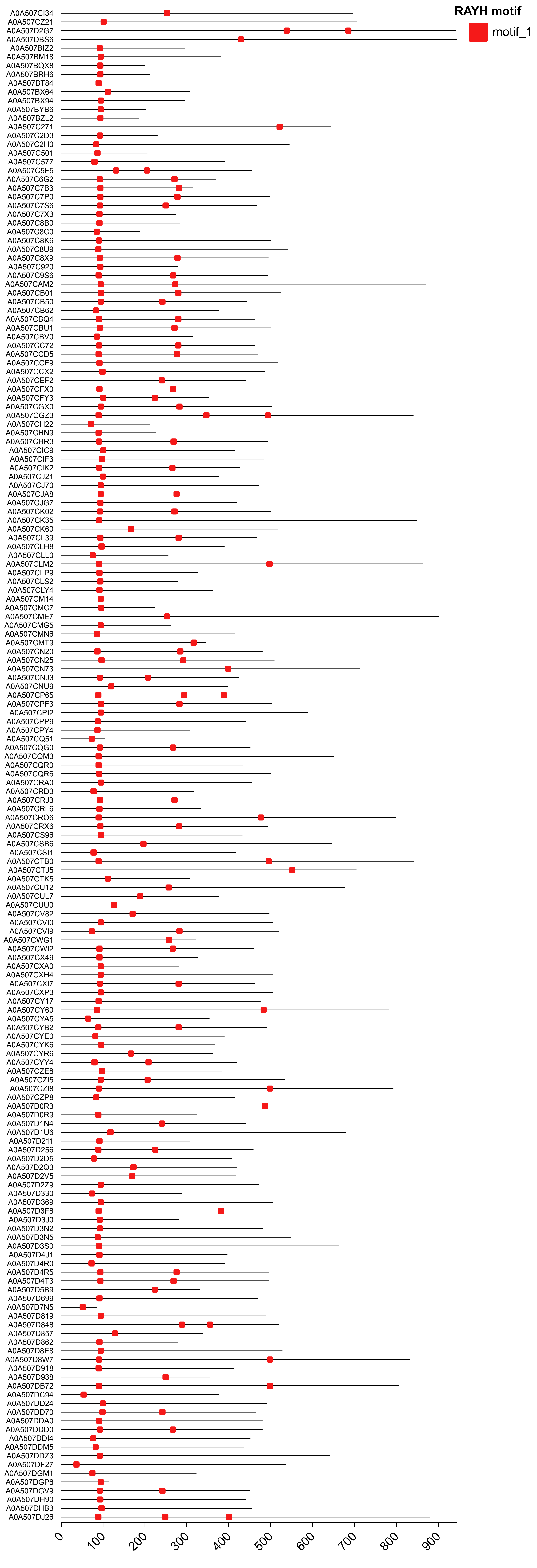


Location of RAYH motifs identified by MEME scan
