## Supplementary material for "Structure-guided secretome analysis of gall-forming microbes offers insights into effector diversity and evolution": Fig. S4

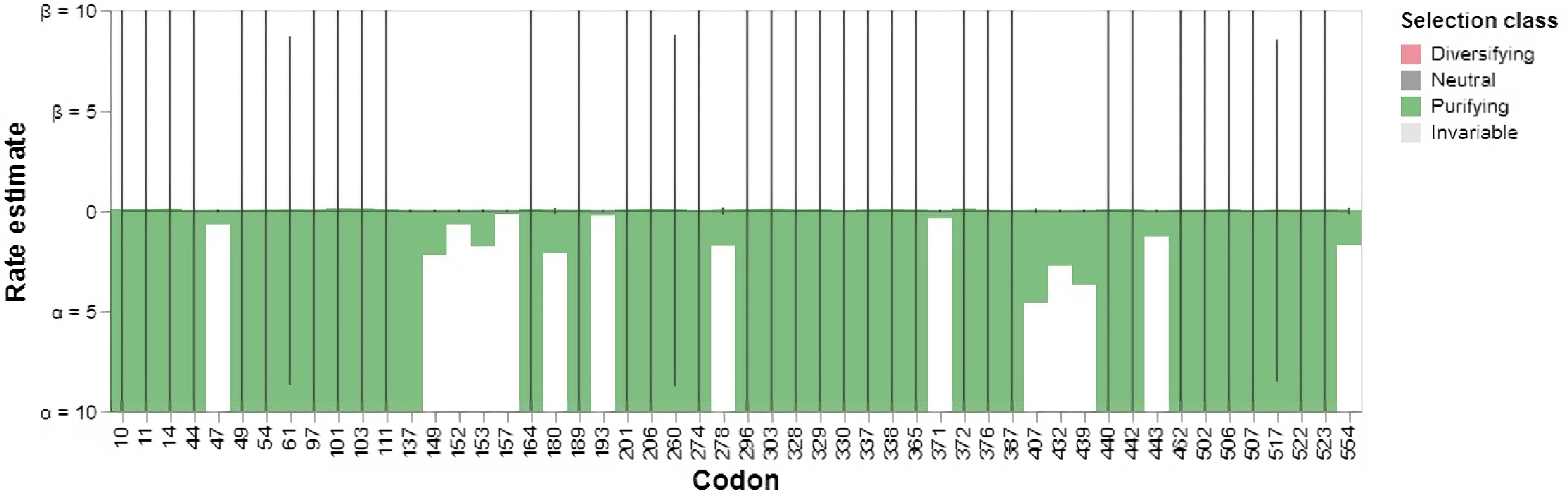


| partition | codon | alpha | beta | alpha=beta | LRT | p-value | Total branch length | class |
|---|---|---|---|---|---|---|---|---|
| 1 | 10 | 10000.000 | 0.101 | 22.752 | 9.270 | 0.0023 | 1236.211 | Purifying |
| 1 | 11 | 7142.863 | 0.106 | 23.172 | 8.367 | 0.0038 | 1259.052 | Purifying |
| 1 | 14 | 848.039 | 0.116 | 22.832 | 8.690 | 0.0032 | 1240.585 | Purifying |
| 1 | 44 | 10000.000 | 0.028 | 22.960 | 46.852 | 0.0000 | 1247.529 | Purifying |
| 1 | 47 | 0.681 | 0.039 | 0.073 | 8.328 | 0.0039 | 3.971 | Purifying |
| 1 | 49 | 346.757 | 0.050 | 22.991 | 20.648 | 0.0000 | 1249.187 | Purifying |
| 1 | 54 | 210.704 | 0.072 | 16.600 | 11.766 | 0.0006 | 901.957 | Purifying |
| 1 | 61 | 10000.000 | 0.091 | 8.700 | 11.740 | 0.0006 | 472.695 | Purifying |
| 1 | 97 | 10000.000 | 0.073 | 22.752 | 13.964 | 0.0002 | 1236.211 | Purifying |
| 1 | 101 | 10000.000 | 0.138 | 22.960 | 7.459 | 0.0063 | 1247.529 | Purifying |
| 1 | 103 | 367.665 | 0.129 | 22.975 | 7.222 | 0.0072 | 1248.351 | Purifying |
| 1 | 111 | 2588.108 | 0.090 | 23.179 | 7.352 | 0.0067 | 1259.438 | Purifying |
| 1 | 137 | 20.256 | 0.030 | 0.068 | 9.884 | 0.0017 | 3.715 | Purifying |
| 1 | 149 | 2.212 | 0.020 | 0.055 | 10.389 | 0.0013 | 2.988 | Purifying |
| 1 | 152 | 0.680 | 0.021 | 0.049 | 7.612 | 0.0058 | 2.637 | Purifying |
| 1 | 153 | 1.769 | 0.040 | 0.093 | 6.793 | 0.0091 | 5.042 | Purifying |
| 1 | 157 | 0.168 | 0.012 | 0.028 | 6.741 | 0.0094 | 1.545 | Purifying |
| 1 | 164 | 2453.442 | 0.103 | 1588.494 | 9.040 | 0.0026 | 86310.509 | Purifying |
| 1 | 180 | 2.103 | 0.063 | 0.128 | 8.398 | 0.0038 | 6.966 | Purifying |
| 1 | 189 | 10000.000 | 0.057 | 23.172 | 19.607 | 0.0000 | 1259.052 | Purifying |
| 1 | 193 | 0.227 | 0.010 | 0.030 | 7.109 | 0.0077 | 1.623 | Purifying |
| 1 | 201 | 631.706 | 0.091 | 24.247 | 11.708 | 0.0006 | 1317.443 | Purifying |
| 1 | 206 | 1310.306 | 0.110 | 22.794 | 8.489 | 0.0036 | 1238.486 | Purifying |
| 1 | 260 | 995.306 | 0.100 | 8.726 | 7.996 | 0.0047 | 474.150 | Purifying |
| 1 | 274 | 83.596 | 0.000 | 11.178 | 72.682 | 0.0000 | 607.342 | Purifying |
| 1 | 278 | 1.730 | 0.071 | 0.183 | 6.887 | 0.0087 | 9.958 | Purifying |
| 1 | 296 | 10000.000 | 0.103 | 22.752 | 7.584 | 0.0059 | 1236.211 | Purifying |
| 1 | 303 | 10000.000 | 0.117 | 22.960 | 9.730 | 0.0018 | 1247.529 | Purifying |
| 1 | 328 | 3422.483 | 0.087 | 23.172 | 10.425 | 0.0012 | 1259.052 | Purifying |
| 1 | 329 | 10000.000 | 0.102 | 22.752 | 6.670 | 0.0098 | 1236.211 | Purifying |
| 1 | 330 | 7477.059 | 0.019 | 23.172 | 53.140 | 0.0000 | 1259.052 | Purifying |
| 1 | 337 | 9961.991 | 0.090 | 22.752 | 10.156 | 0.0014 | 1236.211 | Purifying |
| 1 | 338 | 10000.000 | 0.106 | 22.752 | 9.411 | 0.0022 | 1236.211 | Purifying |
| 1 | 365 | 210.519 | 0.078 | 1717.485 | 10.301 | 0.0013 | 93319.206 | Purifying |
| 1 | 371 | 0.360 | 0.018 | 0.043 | 7.542 | 0.0060 | 2.312 | Purifying |
| 1 | 372 | 7729.926 | 0.123 | 22.752 | 9.124 | 0.0025 | 1236.211 | Purifying |
| 1 | 376 | 10000.000 | 0.058 | 22.752 | 17.254 | 0.0000 | 1236.211 | Purifying |
| 1 | 387 | 158.670 | 0.010 | 14.985 | 48.804 | 0.0000 | 814.201 | Purifying |
| 1 | 407 | 4.599 | 0.036 | 0.110 | 8.027 | 0.0046 | 5.996 | Purifying |
| 1 | 432 | 2.737 | 0.009 | 0.045 | 16.126 | 0.0001 | 2.472 | Purifying |
| 1 | 439 | 3.690 | 0.023 | 0.061 | 11.810 | 0.0006 | 3.310 | Purifying |
| 1 | 440 | 956.920 | 0.096 | 22.820 | 7.999 | 0.0047 | 1239.908 | Purifying |
| 1 | 442 | 1696.780 | 0.092 | 22.777 | 15.662 | 0.0001 | 1237.608 | Purifying |
| 1 | 443 | 1.281 | 0.000 | 0.037 | 24.456 | 0.0000 | 2.024 | Purifying |
| 1 | 462 | 10000.000 | 0.053 | 22.960 | 21.250 | 0.0000 | 1247.529 | Purifying |
| 1 | 502 | 10000.000 | 0.069 | 22.752 | 11.054 | 0.0009 | 1236.211 | Purifying |
| 1 | 506 | 1295.665 | 0.091 | 22.794 | 12.796 | 0.0003 | 1238.529 | Purifying |
| 1 | 507 | 10000.000 | 0.025 | 22.752 | 27.078 | 0.0000 | 1236.211 | Purifying |
| 1 | 517 | 53.890 | 0.075 | 8.516 | 15.285 | 0.0001 | 462.738 | Purifying |
| 1 | 522 | 8851.167 | 0.063 | 309.454 | 19.355 | 0.0000 | 16814.128 | Purifying |
| 1 | 523 | 10000.000 | 0.082 | 22.752 | 12.002 | 0.0005 | 1236.211 | Purifying |
| 1 | 554 | 1.710 | 0.053 | 0.135 | 7.049 | 0.0079 | 7.338 | Purifying |

Codons under purifying selection in the CCG effector family. Plots have been downloaded from the Datamonkey server, which hosts the FEL package. The indicated codons are numbered according to the codon-aligned nucleotide alignment file, where positions with 50% gaps have been trimmed.
