## Supplementary material for "Structure-guided secretome analysis of gall-forming microbes offers insights into effector diversity and evolution": Fig. S5

```
            1       10              20                           30
A0A024FWW3  MHHLRPCALFVTTL...LLL...KSF.........SLITSLS.G.........ILNVTLS
A0A024FUJ7  .MVLR..SSFIPILMELLIIN..AGA......SKIGRLQSAS.S.........YSRLLAV
A0A024FVG4  ...MWSN.....IV.LFLLAAYHQ....VSQASGYRI................STILP.D
A0A024GFU0  MVHKASPL....TL.VFLIGSL........GCCKSTRI...............TFTTPKR
A0A024FV15  ..MFRQNW..........................LFFSVVWLLNTVTDANNQILITATLT
A0A024FW891 MKVTCVTR....IAVLY...H.............FYASCAAQLGS..........MKITV
A0A024FTL6  MKSLPLTT....ILVSLLARHLHGSFG..NFVEESYLMAMMDNGY..........TNMER
A0A024GNE5  ..MVRSTR....SLISLVIVTITSAHLVEGSFTKLY...................GEFPY
A0A024FWW7  MGYFRAIT....VLATFVSCHSSPSQY..DVQTKPYDFKIQEQ............NS.AK
A0A024FWT1  ...MRVQQ....ILSVLLIHS..TGYQ..TAQSSPYMRTIIVS............TTFNV
A0A024GUG61 ...MPDAR....ILLAFLLCCVLCAFQ..RACCDLKT.QTWTL............ISDDV

               40        50        60        70
A0A024FWW3  ETEDFDACHNCLISSLGIKQIHLIAAARAFRLYWV................EGHSDTMTA
A0A024FUJ7  TTAQVEDCHTCLLRNVGVERITLVASNKITRIYLI................KSRVTVLES
A0A024FVG4  KYTRVRKCRRCLQRLIP.NAIFSYGYG.VGYYQRSI....RQNVLLMYADITDDIPYRS.
A0A024GFU0  NSMRVLKCRDCILSNFYADEITLISEG.AHNVYSSL....ARTKFMYHANIPDDFS....
A0A024FV15  HNSDLGKCQSCLTDIAGMDRLSLALPPTDDP........KSLTTTVHFYASGYSASFMR.
A0A024FW891 QTAKYEPCQQCLLEEAGAVHVEEYEGARRGG.........TSYFLL.....G.LIDYIKT
A0A024FTL6  LVHKAAKCQDCMINRAEVTVVSFDA...............ITPFLAKFTIRASRLGFVR.
A0A024GNE5  STSTLKKCQTCLLEQAGARRLILMKKTITGTI.......AQKTVNFEYQTQGPSYFYSN.
A0A024FWW7  LRDAFRNCQQCLLDFAGIVSLRVTEPSSDK...............FVVYITASLMAFIR.
A0A024FWT1  GDTNLLICQHCLLSTGGATRAKLVKFSKPPY........LSGSTHADFEVTASEFGFVT.
A0A024GUG61 KLPEMCSCQNCILNVAGAMRATIIHMDKAKNAYKNVKYSVTAYKNVKYSVTASDFAFMS.

            80       90       100                        110     120
A0A024FWW3  FLSYCSNRVACVLSDQLQQTKL.............PANEM....KATRYRTQQNELESVH
A0A024FUJ7  FLNQCNNEKSCQLQTDIDMFPPWSSI....LWR.NPVLEV....AISRFSMHINDIT...
A0A024FVG4  VVRSCQPKKAQGCYRISDCEEGGIRY..EKLMDVIKKFKMMQVGGPIRLFG.........
A0A024GFU0  ............................................................
A0A024FV15  AKLDCQSDLTCGFITMENTKITDNTI..................................
A0A024FW891 NLIKCSATYQCRGYKGE...PIVFEN.KEE...............EQLIFGHINEKQ...
A0A024FTL6  IQMYCNFNEQCSYLQRLRQRELGSSTPTSTMYNTNPTY....LHECTELLTHLNSKL...
A0A024GNE5  VETYCVGRQYCGILAIGATQPY.HEA.KEAT.............................
A0A024FWW7  VHTICFEEDVCDKMQLSEGSDSANEI.TESTWQT..........................
A0A024FWT1  IEQICRIRTICGPLEVHRSEYKNDQP.I....V...........................
A0A024GUG61 IHSKCVLDGYCSAFELVHSSTPSGFS.TESS.NL..........................

                 130       140       150       160       170       180
A0A024FWW3  SLASSMHEAIVKWSDGSTTSLTESIYEVSSLIWKESSEIPMFFWDDELIRDPLLLKHY..
A0A024FUJ7  ............LDPSRDKKVV......S..NLESYRGIAFFFFDEI.QKMHI.RRHRTV
A0A024FVG4  .................ADLAS...........DESRGESSGAA..KRARLPDLEVNYNT
A0A024GFU0  .................ATILR...........DVCKGTDSCDS....ISFPE.......
A0A024FV15  ............................................................
A0A024FW891 ..............LASNS.................VG......SQTGGKIPLL......
A0A024FTL6  ..............PCKNDKLT...........ELMKGSTLYDTARPNDSLDIL......
A0A024GNE5  ............................................................
A0A024FWW7  ............................................................
A0A024FWT1  ............................................................
A0A024GUG61 ............................................................

                        190       200       210       220       230
A0A024FWW3  KRKEP.....KILIMSYPAVLNSKMPVDDHVNTQYGKTVESLMRCMILPFTPQEAVCREC
A0A024FUJ7  KTRNAQLERKQIDKSTYQHSSVGNILIAQ...DPAVRESPSLARCFYS..YSRRFQCERC
A0A024FVG4  EKR.......QT..........GDASTSS..ANAEHGIEQSKTHCLLFEKNLPEFLFLFC
A0A024GFU0  .PA.......QI..........GVQSV....TNMAPPTHTSTAQCIAVDYRYLT.EVTTC
A0A024FV15  ...............................AADMLG..LSEAVCVL.TDVQTSFECGTC
A0A024FW891 ...........L..........SNDYASQ..TTSMISGRVAKPVCLYRPDKFHSSLCETC
A0A024FTL6  .........................LQSS..DSSMLGQSNGKSLCIFQLHRLVTADCKIC
A0A024GNE5  ...............................EAKPEQSYDKRYQCLVRFSLAHSLSCFRT
A0A024FWW7  ...............................PRTIQIEDPYKIVCITHEDKYVREKCRVC
A0A024FWT1  ..................................LQSRVSRSILCLV..SHATTTKCRDC
A0A024GUG61 ...............................VATEQEAAKSEPMCFILGPQKTTARNRLC

              240        250       260       270       280         290
A0A024FWW3  MAMHSKW.FAVRSIHLYHTSKKRSADVCVFHSNYSSSLIVANCDKV.FC.PAIKILNS.I
A0A024FUJ7  LALHSIS.LGVRAYPFKKL.KKGLDFVFAFASFHTHGKIQENCAKK.YC.LSLNSLSS.A
A0A024FVG4  LQCLEIG.EKEERYLISNPHESDRQWVVRYTSSETPEGTMQNCGSL..C.GNIKWGSS.A
A0A024GFU0  QECLSNHGSYMKKYWV..IIETVYHWMFWYYSS...GQHLLRCTQK..C.GGMKILSR.A
A0A024FV15  LRMVSGS.HTIQKFQLIAKNENE.QYLHVLYT..HQRDIFAPCSESGVCRGQQYEFGY.G
A0A024FW891 FKQKSDT.RSMEKYAMYNMGISS...VYVIYTSDT.NVFLRSCRNLRMCRGKTVVFSE.N
A0A024FTL6  IRQTSGS.KSIVEIQFVLNVE..HYIVYLLRSSKHMEKILKDCSRCSFLQGELSRYH...
A0A024GNE5  LQELSGS.ELMGDFLLRSPLYHHDLHLFLLFTSKNLKEMTEDYFPGYACNGHSVVLSS.D
A0A024FWW7  LREASDS.ESIGEFSFFQDGF..FQFMYVLRTMRTGDDMAEACAAKYKCLHRVENGNNDG
A0A024FWT1  LKEASDS.PFLGNFRTSSVQLRNLFTLHILFTAKEAEYIVQSCSASLACIGTMVELQG.N
A0A024GUG61 LKEVSGS.ELLNEYPVASSYDRITRTIVSLYPKNHK...TDPKLLLLACMGFG.DLKV.D
```

```
              300       310       320        330       340
A0A024FWW3    DCHRLHDIFFISHPNADDKNKLQQEELQEVLTTIAGN.RMARRHKVDGVFQEAVASLNS.
A0A024FUJ7    ACDNLDRMFPQDKKTLHDDGTSANGAENQ.IEIVPPNHSQASISSQEDIYFRYIASDLDF
A0A024FVG4    HCDAVHIDYTT.................TLSDLPLDSI..SILE...............
A0A024GFU0    YCEPHMIDLNE.................LAEKQPYNKV.....R...............
A0A024FV15    TCSNLARKLYMRQFS............PFSGA...............TLKFN........
A0A024FW891   LCTLFRHKISSRKLQQE.......TA.....SMLPQVDT..II.................
A0A024FTL6    ..EYLCDETYT...............................................
A0A024GNE5    ACDSFVERIFPKS....................DDDQ..RLQRSPSPKRAK....AEF
A0A024FWW7    ACDAMHNTGVSK...........................HYKSKGITV.........
A0A024FWT1    VCRFLVPENVS...................PDI..........TWKV.........
A0A024GUG61   PSNYVSPDSCQNLIMKT......LVFSPLTIKKPLISR..SMFKTDSPST.........

                                      350       360        370
A0A024FWW3    ........................FNILHAKEYVCIKVSTQP......ILQWQCISCVLK
A0A024FUJ7    GSDLGTELDTPTKNVKLGIEIQVIEQFLSQTEISCMRVRRDQ......NDRLFFLRCLSE
A0A024FVG4    ...................................:........VKYPKLNKQDCLRCVSF
A0A024GFU0    ...................................:........VAFSKTNRAKCIQSLAE
A0A024FV15    ..AAEVPFDV.........MIPFIYETQHVSTRCFLASTTL......KLRTRCLRCMAL
A0A024FW891   ..GNIKIWPPTL.SKSIEIEKSEAELRAILFQSYCISTTASY......IGNHNCWAGLQK
A0A024FTL6    ....GVPRQE...............ISEYPDMMCVQVTLTPSLFYSNDDMLQCMRYLAA
A0A024GNE5    LQRYGLSMRKEIRANHL..EIVGQIEETLGMEYHCFRTTLAA.....G.KSGSCLSHLSD
A0A024FWW7    ..TNADMNANDKNERKL..QCVAAE....LLEH.TSSATIHY.....YLSMQECMECLAH
A0A024FWT1    ..TNGLPWSGPTGTDGF..KNLQAKYQSGTLKHLCLKVPKQG.....LQAATRCMYLLAE
A0A024GUG61   ..SSGNSWERAIQLEGI...............PLCILVTFAP.....GSKVSECMTTFFE

          380            390                         400              410
A0A024FWW3    DVKGYRIVIF....SS............LYSTLIF..TDTIK...PMS.....SCFKC..
A0A024FUJ7    S.AGNDNPIFLVRSSS............LTELILFF.KSSLLKVDFAR.....NCFEEYG
A0A024FVG4    NLHAIN......VHNGD..........DDMHSIFAATTQAVQLKWND......LLKTCK
A0A024GFU0    ATNILT..RLGHSDSGE...........EDSSALIFIGSGTIP..IN.......NLSGCR
A0A024FV15    ESSGIA..VLVDDTSFAH.......EHEITSELYVL....LISEKYQNKKEIHGCEQVCN
A0A024FW891   QSGGMV..VLDRIETFR........TPEKQGEFLLY....LFTNGPPDA..VLANIQYCG
A0A024FTL6    ADGNIA..YWMAIGAVPS..........LPAYFLVS....YVPMEFNNV..GETQVQTCG
A0A024GNE5    TIGKTT..LFMGRSFKDRTGSTNEQNGSQKLYFYYI....LLPAKSGDQ..LFRSSGACG
A0A024FWW7    KTNSPV..HRMRISAE.............HPREMYF....LFIMEADPQ....DSSDCGT
A0A024FWT1    RIDGLV..IMLHKGL...............KEEHVN....IIGFQVDSS..SADSITVCG
A0A024GUG61   NFNGEA..VFLGKKL...............EYHYFY....LFGSKKNKS..VFSFKKPCA

                         420       430       440                        450
A0A024FWW3    ..RMIEVLHP.DDCRQNLAHHSNNLEVYPKPS...........NFEMH.....NRLDRAC
A0A024FUJ7    GSDIVPLLNPPSFCDYRIKYKIKSNTYPELR...........TLTKT.....KPQTELC
A0A024FVG4    TS.........KKCSGVKDKSSQISSSLSEFR....SIKPLSHKGGDSFSGRETTPTYLC
A0A024GFU0    TL.........ESIETAHDH..........T....EMEPGPHPFIMSYPGINVEPS.QC
A0A024FV15    K....PSRKNKEACSKASLQAIDHSSPSGKIYYNVALYPK..LNQIDPSPLRQP.LTSNC
A0A024FW891   EAK.......FSDHCM.LDAVHVDTLTPTHDIFEIMERKKP..GQQGKLAPN...LHGECF
A0A024FTL6    SDRTIIEMSQIGNCFMNYYNQFEPSPPFETQIS...THHP..HPDTRFPPS.TLTTSTVC
A0A024GNE5    A...VEYAQSKDECSSLLNQHLFVSPQYAPKAK....RFKS..LSAKPT..SPSTQQPSLC
A0A024FWW7    C...TIKKVETKECYIEYERKFDTLPIYDQIS....VNMD..LTLRRK..NVLFHPTSCF
A0A024FWT1    K...KMLTVDQTICNELNAHQFDTTPISNQIS....VLKS..QKAQMEQANVKIEYTSEC
A0A024GUG61   ....TIKKADLEECKAFHSHHYDTSSVQRQIL....SWKT..QESEKVILHGFPSHQTRC

                                      460       470              480
A0A024FWW3    ................FTVYHTKSDARKCPSCIKSFLGITNEA.......QLIRQSPLL
A0A024FUJ7    ................VLVSHITSKYKSCGRCVKTFFDPMYKG........WVSMP....
A0A024FVG4    MHYT...............SRGVLPADTCKECIKEIIDSKHSRSQKGETRIMSRKSGLI
A0A024GFU0    LHVIRPQSPWLSPKSDFTTFRVEDIENAECENCIRTLEPGYPI....SGVRLTTYMSSFL
A0A024FV15    ................IWVQHELSNEVTCQTCLKENAGHSVE........KLSVVDFLI
A0A024FW891   ................LLQH.LYDDESKCRAFFPESWKH...........KLSAIAFLV
A0A024FTL6    ................LVAH.YETK....SRFKRSF..KTGSM.......ELSAHSSLI
A0A024GNE5    ................IWIF.SVTP.QDCLIWLHERLDTSASLT......MLSRTTVLL
A0A024FWW7    ................IVNY.AKRDEDDCMRCIRKIFHSRGSFH....GSRMSAISFIV
A0A024FWT1    ................VWTR.HDPEIGKCTSCLGRLGISEESIL......PLSQSTSLT
A0A024GUG61   ................AVAQ.HRKDAIDCQRCVQHFGNHVIA.N......TMTDSTFLV

          490            500       510                520       530
A0A024FWW3    FDISMSKWKA.....TPNKFRNLLSTCV.......RKYKCTRIQNVPNIFCMHHRDFIMA
A0A024FUJ7    .PTHSSLWKDSKERNPSTMVDNVLLHCI.......RTKRCSKIQLYPKMVCKMHEEIMEL
A0A024FVG4    .......F......LTEKLPVQKVA.........LQCDEFCDWVSGLTEDVCRLLQNIETM
A0A024GFU0    .......F......LQSAPHRFYIA........DNCKK.CLSITVFPNSFCGQPE.....
A0A024FV15    .......Y......REKNLAMDSIYKCIST....FKRSNLCKKMVSAPDQFCLYYKSLRDG
A0A024FW891   ...S...F......YADTNGHFNLKS.......SLFRDTHCKGVDLMPDAICLYELDRIKF
A0A024FTL6    ...E...L......QAT.....SMAYPMH....GILSIEFDKLVFIPQEMCVYERPKLAS
A0A024GNE5    ...H...T......VDSSVAES..........VHLPSSCHVLHDIPMSVCTIEEELTRR
A0A024FWW7    .......R......VDSSEIQRDLPYCITK....SHERACMMISYVNNRICLGNEAEYL.
A0A024FWT1    .......L......LATNFEKAKLASCINAGNKKRTDLQCEHFVFVPNFMCEYDLTQPEQ
A0A024GUG61   .......S......LPDKFDESILATCIKDVYYRNDATKCSHFVFLPREMCEYARMKTSA
```

```
              540       550         560       570                 580
A0A024FWW3    RLTEQSKADTFTHR..LAPVGAWPSLSMDLFTPPRRYSP..........HSSPAYSSQIW
A0A024FUJ7    HVKGNVINEKSMSKFDRMTDRVWPSADPHLSG.........................SEN
A0A024FVG4    N....S........GDSQVPGQ..KVEEYHFVTAITY....LS...TKPVTSHLWYLQ..
A0A024GFU0    T....S........KDPPTP......QEYLV.............................
A0A024FV15    SRSSI...........VPYPEN..RI..................................
A0A024FW891   KSDLNLR............PQELTST............SGSDHSE.......ASFA....
A0A024FTL6    PKVINRADRKS...NDATKPTSISSRGKSEF........GSSHS....PSNAKSFPTISG
A0A024GNE5    QHSHRSV.........DPNPHPFPEADEDSMSTLKQQPAMSSTSKMTAQSSSPSFGSIIT
A0A024FWW7    .RPVQE......................................................
A0A024FWT1    TFSSQEQQ......QGTRYPVAFDTVGSSS..NPRNFSSGYLASSISHQQSLQLFGKQIK
A0A024GUG61   KSPSGASS......SN............................................

                590                600       610       620       630
A0A024FWW3    PVSGDIALVVFTYA.......DFELYHCLHCLAEHTEIFFFYRNEVAHNGYVWMRPESST
A0A024FUJ7    ..LLDRPALMIVYD.......AIDPKECLRCLILCTQVFYSYKFKGNRRGYIWMHQSSSD
A0A024FVG4    ...........FKQ..GEGDMEEAFKDLAFCIAVVSNAYVVS....VRLRYLIMSPFDYK
A0A024GFU0    ...........VTI..HQTEFDYMFDHIVAHVAFHTTVYLVD....MNCRSILTSAGSRQ
A0A024FV15    ......GNVGI......GMP.SKPNSKCFTCLTSILDILII......DQNHFWMLGAPRE
A0A024FW891   .....EAKVTLYKWTDKGMDNEDGMKDCFQCLTRKRNIVVLS....VTNQYVWMVEYKMD
A0A024FTL6    R....DDAIVLVSL.......IPENPQCLACLALSNEVLLVS....TVKNYVLMFASQHD
A0A024GNE5    V...IHLNKDVNAL.......PLNLFNCATCLIIYEEVLLIS....VELAHVLVTKEDRD
A0A024FWW7    ............................................................
A0A024FWT1    L...EPPVTMVSMK.......APKASDCAASLALEYTVLMIS....IKKQYVWMLYDREE
A0A024GUG61   Q...AQPNVMVVTW.......EVLNIGCLQDIALYHDVLLVS....KTDRYLWMLRVTSD

              640                        650       660       670        680
A0A024FWW3    VLTVCLG.KY.....CQS..........LSHGESVLRTRRPFGLRQLTNVDIKSSV.RFS
A0A024FUJ7    LLSPCISGNS.....CTE..........LWHDETAEQLAISDDMRPWSVVDIGDLN.DF.
A0A024FVG4    HVA.TICNIPDIIRITHFDN..............VYSGI...FPQPLSLQSITSSL.GIS
A0A024GFU0    TIA.TICASAARS..VGYPDV............CVYGTVQANRLQPMAFSLIPQLF.GLP
A0A024FV15    GDCVSECGDI...KMVVRLE..................LDVSSIRPMSSGSIEGRFAKER
A0A024FW891   FPNCPSCADK...V..................KPGPQIPYDHNLRQILARDVNLLF.GIG
A0A024FTL6    DWH.CGCEFV...SFTQFDYVKTHKWRQLSSIEIVYHLDDPQFNKRIMGENVGLLL.NEL
A0A024GNE5    VNGFTGCEQV...CSFSE..........NSKPEPVSHIEDFRLLHPLSMDDLVSIF.CGD
A0A024FWW7    ............................................................
A0A024FWT1    PLSAEGCTNL...HMCHR..........L........QQDVHMLRAVSAEDVIRNL....
A0A024GUG61   ELR.KRCKEV...RIEKE..........Y........FFDVHFLRSISAFDMKKFI.GPT

A0A024FWW3    PGLQ........................................................
A0A024FUJ7    ............................................................
A0A024FVG4    GHTGPKIFISDRLEQ.............................................
A0A024GFU0    QAIDLNLAIIN.................................................
A0A024FV15    YSLRYEKMLDNGHL.........YEYAEML..............................
A0A024FW891   EVKKSRKRSK..................................................
A0A024FTL6    YPKKSNRYI...................................................
A0A024GNE5    IGEKKRRYVDRGNKRKDQINISYVDYHKCLNYLTTHPLDVAIPKEMPTSSDKVTETRTKS
A0A024FWW7    ............................................................
A0A024FWT1    ............................................................
A0A024GUG61   ............................................................

A0A024FWW3    ..............
A0A024FUJ7    ..............
A0A024FVG4    ..............
A0A024GFU0    ..............
A0A024FV15    ..............
A0A024FW891   ..............
A0A024FTL6    ..............
A0A024GNE5    KNKRKMSSGDERSK
A0A024FWW7    ..............
A0A024FWT1    ..............
A0A024GUG61   ..............
```

Multiple alignment of representative members of CCG sequence-based clusters. Conserved residues (>70% similar by biochemical properties in ESPript3.0) are highlighted.
