## Supplementary material for "Structure-guided secretome analysis of gall-forming microbes offers insights into effector diversity and evolution": Fig. S6

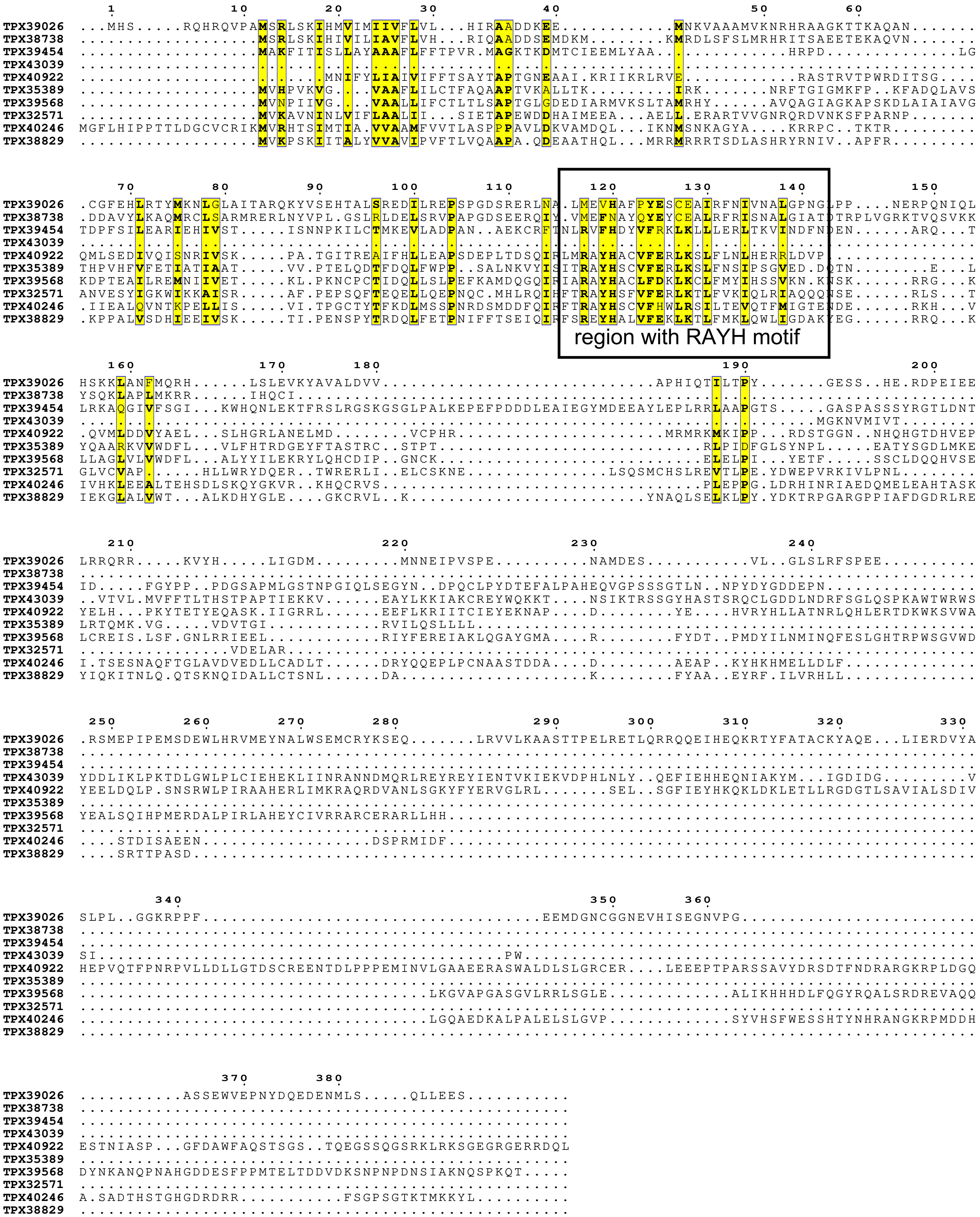


Multiple alignment of representative members of AvrSen1-like sequence-based clusters. Conserved residues (>70% similar by biochemical properties in ESPript3.0) are highlighted.
